# A rapid, sensitive, scalable method for Precision Run-On sequencing (PRO-seq)

**DOI:** 10.1101/2020.05.18.102277

**Authors:** Julius Judd, Luke A. Wojenski, Lauren M. Wainman, Nathaniel D. Tippens, Edward J. Rice, Alexis Dziubek, Geno J. Villafano, Erin M. Wissink, Philip Versluis, Lina Bagepalli, Sagar R. Shah, Dig B. Mahat, Jacob M. Tome, Charles G. Danko, John T. Lis, Leighton J. Core

## Abstract

Tracking active transcription with the nuclear run-on (NRO) assays has been instrumental in uncovering mechanisms of gene regulation. The coupling of NROs with high-throughput sequencing has facilitated the discovery of previously unannotated or undetectable RNA classes genome-wide. Precision run-on sequencing (PRO-seq) is a run-on variant that maps polymerase active sites with nucleotide or near-nucleotide resolution. One main drawback to this and many other nascent RNA detection methods is the somewhat intimidating multi-day workflow associated with creating the libraries suitable for high-throughput sequencing. Here, we present an improved PRO-seq protocol where many of the enzymatic steps are carried out while the biotinylated NRO RNA remains bound to streptavidin-coated magnetic beads. These adaptations reduce time, sample loss and RNA degradation, and we demonstrate that the resulting libraries are of the same quality as libraries generated using the original published protocol. The assay is also more sensitive which permits reproducible, high-quality libraries from 10^4^–10^5^ cells instead of 10^6^–10^7^. Altogether, the improved protocol is more tractable allows for nascent RNA profiling from small samples, such as rare samples or FACS sorted cell populations.

## 1. Introduction

Next-generation sequencing technologies are now routinely used to measure gene expression levels in a highly sensitive and comprehensive fashion. RNA-seq facilitate simultaneous detection, identification, and annotation of many classes of cellular RNAs. Traditionally, however, this technology primarily measures steady-state RNA levels, which are a consequence of equilibrium between RNA transcription, processing, and degradation. As a result, many unstable RNAs, particularly eRNA and some lncRNAs, are not easily detected with these approaches. Alternatively, ChIP-seq allows for identification and quantification of RNA polymerase II (Pol II) associated DNA. This produces a genome-wide map of both transcriptionally active and inactive Pol II without strand specificity, thus the direction and transcriptional status of the polymerase are ambiguous. Furthermore, the relatively high background in ChIP-seq as compared to RNA-based methods obfuscates comprehensive transcript and regulatory element detection. To address these shortcomings, various methods of measuring transcription by genome-wide profiling of nascent RNA have now been developed including GRO-seq (Core et al., 2008), PRO-seq (Kwak et al., 2013), NET-seq (Churchman and Weissman, 2011), and TT-seq (Schwalb et al., 2016). Characteristics of these assays are reviewed in (Wissink et al., 2019).

GRO-seq and PRO-seq are modern, genome-wide improvements of the nuclear run-on assay, which has been in use for approximately 60 years (Weiss and Gladstone, 1959). Over the years, run-on assays have been instrumental in the study or discovery of various forms of gene regulation including, steady state transcription levels and mRNA turnover (Derman et al., 1981; Powell et al., 1984), promoter-proximal pausing (Gariglio et al., 1981; Rougvie and Lis, 1988), transcription rates (Bentley and Groudine, 1986; Hirayoshi and Lis, 1999; O’Brien and Lis, 1993), and 3’-end processing and termination (Birse et al., 1997). The nuclear run-on reaction works by adding labelled nucleotides to polymerases that are halted in the act of transcription yet still transcriptionally competent. The polymerases incorporate the exogenous NTPs and the labelled nascent transcripts can then be detected and quantified. In 2004, the assay was adapted for macro-array by spotting probes of whole yeast genes on to nylon membranes (García-Martínez et al., 2004). In 2008, the nuclear run-on assay was expanded to cover the entire genome in global run-on and sequencing (GRO-seq) (Core et al., 2008). GRO-seq employs a BrU-containing run-on reaction, enabling affinity purification and subsequent high-throughput sequencing of the entire nascent transcriptome. This allows sensitive and strand specific measurement of all transcriptionally engaged RNA polymerases, but resolution is limited because run-on length can only be loosely controlled via limiting nucleotide titration.

To address this shortcoming, precision nuclear run-on and sequencing (PRO-seq) uses a similar run-on and sequencing strategy but substitutes bulky, biotin-tagged NTPs instead of BrU (Kwak et al., 2013; Mahat et al., 2016). Biotin effectively halts eukaryotic RNA polymerases after incorporation of a single NTP, which enables genome-scale profiling of transcriptionally engaged polymerases with base-pair precision. PRO-seq and the associated technology ChRO-seq (Chu et al., 2018; Kwak, 2013) have now been used to study the transcriptional kinetics of the heat-shock response in human (Vihervaara et al., 2017), mouse (Mahat et al., 2016), and fly (Duarte et al., 2016), for *de novo* discovery of promoters and enhancers in cells and tissues (Chu et al., 2018; Danko et al., 2015; Kruesi et al., 2013; Kwak, 2013; Wang et al., 2019), for transcription rate detection (Danko et al., 2013; Jonkers et al., 2014), and detection of RNA stability (Blumberg et al., 2019; Core et al., 2014).

The current, published PRO-seq protocol (Mahat et al., 2016) is undoubtedly time consuming, technically challenging, and requires significant amounts of starting material (0.5–2 × 10^7^ cells). This is largely due to multiple streptavidin bead binding and subsequent elution steps (Fig. 1A), which require technical finesse with phenol:chloroform extraction and ethanol precipitation of nucleic acids and present repeated opportunity for loss of material. This has limited the adoption of PRO-seq by inexperienced laboratories and impeded studies of transcription in experimental systems that utilize rare or precious biological samples.

**Figure 1:**
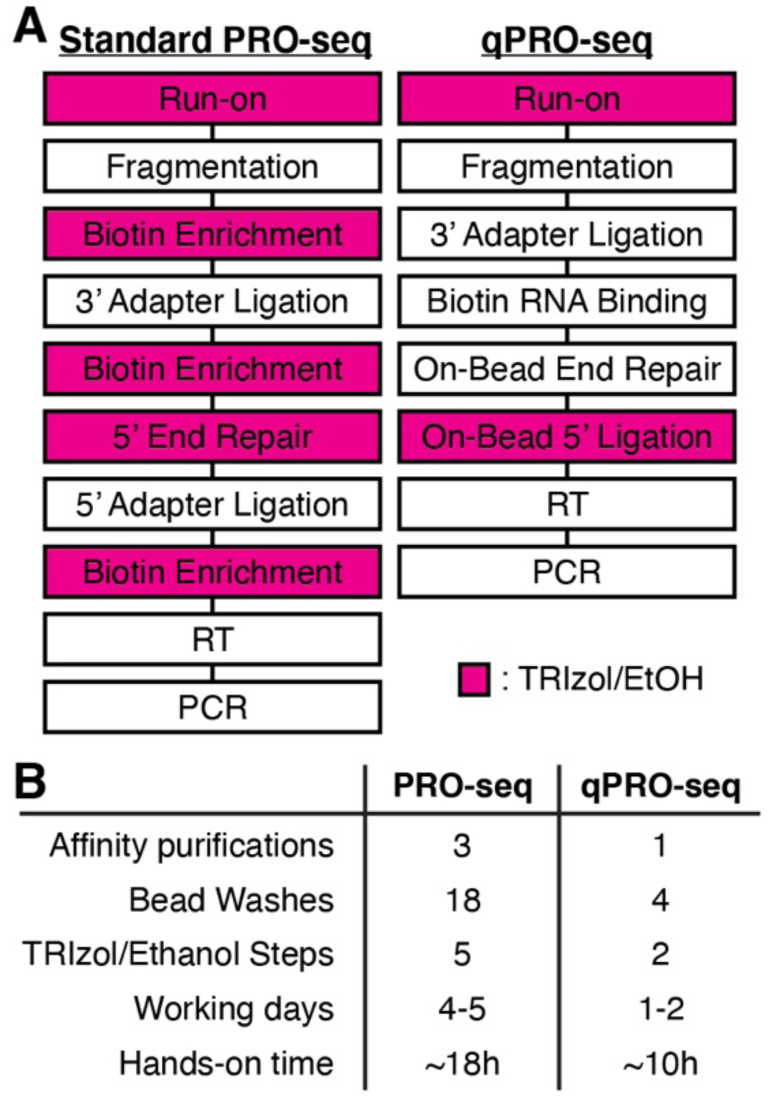
A rapid and efficient PRO-seq protocol. **(A)** Overview of changes to the PRO-seq protocol implemented in qPRO-seq. **(B)** Comparison of steps and time required to complete each protocol.

To address these shortcomings, we optimized the PRO-seq protocol to simplify library preparation and facilitate use of scarce input material in an improved protocol deemed qPRO-seq (<u>q</u>uick <u>P</u>recision <u>R</u>un-<u>O</u>n and <u>seq</u>uencing; Fig. 1). The original protocol requires 4-5 working days to complete, and included three bead binding steps, and five phenol:chlofororm extractions and ethanol precipitations. By performing 3’ adapter ligation to hydrolyzed total RNA instead of affinity purified nascent RNA, we eliminated one bead-binding step. We have validated that this ligation reaction is equally efficient to the standard PRO-seq ligation to purified nascent RNA (Fig. 2A). Downstream enzymatic reactions are then performed while nascent RNA is affixed to streptavidin beads (Fig. 2B), which eliminates another bead-binding step. On-bead reactions are performed in 2X volume to aid in handling, and simple bead washing steps replace numerous phenol:chloroform extractions and ethanol precipitations. The resulting single bead-binding protocol is much faster and easier than the original PRO-seq protocol (Fig. 1). It is feasible to start from permeabilized cells and end with adapter ligated cDNA in a single day (∼10 h; Fig. 1B). Furthermore, an option for column-based purification of RNA after the run-on reaction can eliminate another organic extraction step. Reverse transcription can be performed on beads, which completely eliminates organic extraction from the protocol, albeit with reduced efficiency.

**Figure 2:**
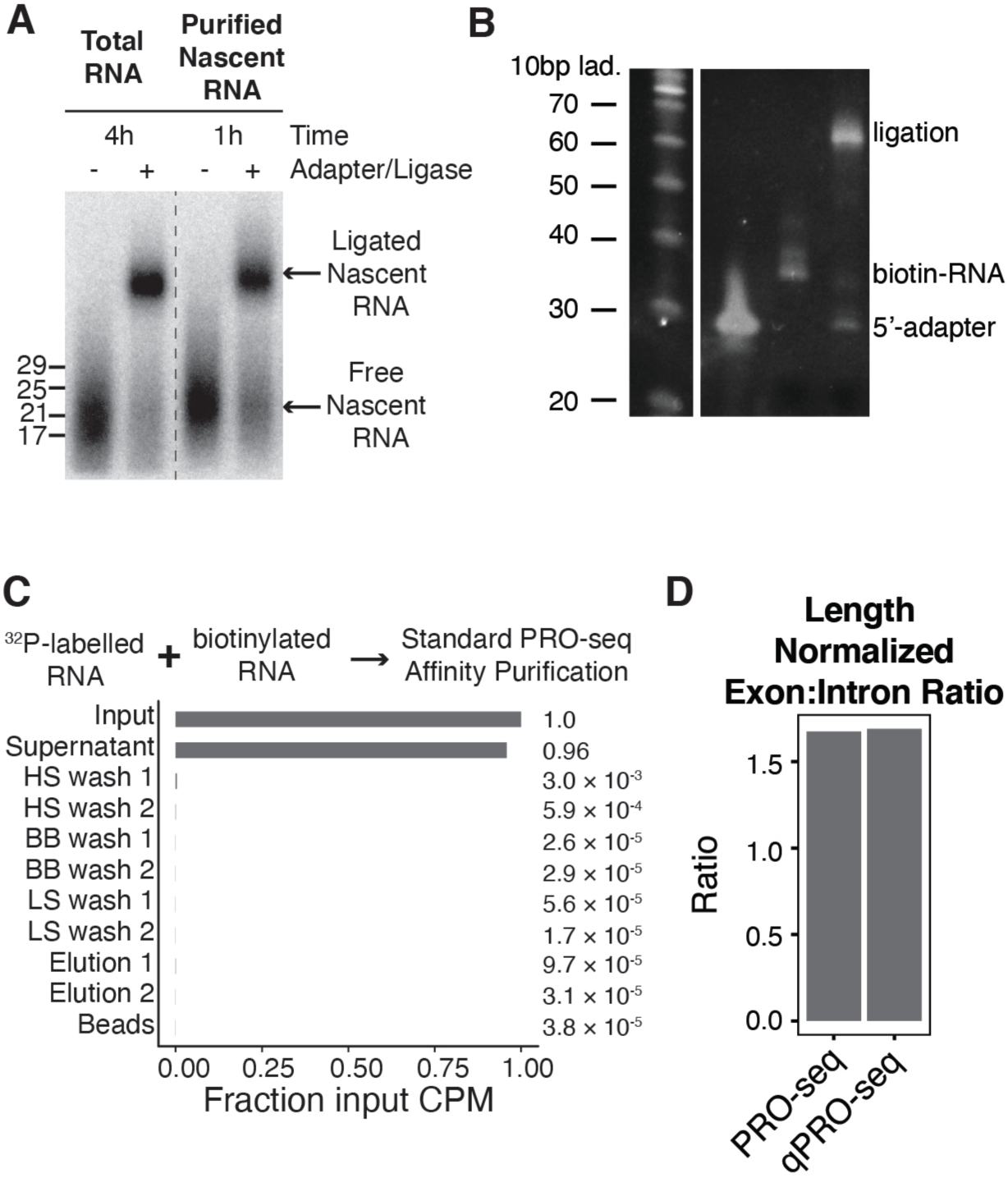
Validation of key enzymatic steps. (**A)** Ligation of the 3’ adapter in hydrolyzed total RNA vs. purified nascent RNA. ^32^P labelled nascent RNA was ligated for 1 h using the ligation conditions in this protocol or using the standard PRO-seq ligation conditions in (Mahat et al., 2016). **(B)** Efficiency of ligation to synthetic biotinylated RNA in 15% PEG8000 **(C)** Stringency of biotin-RNA affinity purification with MyOne C1 Streptavidin beads. Excess ^32^P labelled non-biotinylated RNA was mixed with biotinylated RNA, and CPM of each fraction was assessed using liquid scintillation counting. **(D)** PRO-seq and qPRO-seq show similar levels of exonic reads relative to intronic reads genome-wide.

In theory, reducing the number of affinity purifications and ligating adapters to bulk RNAs could reduce the specificity of the assay for nascent RNA. However, we have found that MyOne Streptavidin C1 beads, which have higher binding capacity per substrate area and are negatively charged to repel non-specific nucleic acid binding, sufficiently enrich nascent RNA over other cellular RNAs (Fig. 2C). Importantly, if nascent RNA was contaminated with mRNA, the number of reads mapping within exons would increase relative to reads mapping within introns. However, when we compared the length-normalized ratio of exonic reads to intronic reads, we observed no detectable difference between the original protocol and the improved protocol presented here (Fig. 2D).

Additional protocol changes have further simplified and improved the protocol. Careful titration of adapters eliminates the need for PAGE purification (Fig. 3A–B), which is difficult and time intensive and can introduce insert-size bias in the final libraries. Incorporation of a dual-UMI strategy reduces concern about PCR-duplicate reads. Furthermore, this eliminates the need for time-consuming test-amplification, except as an initial troubleshooting step when establishing the assay in a new cell type or with a new amount of input material.

**Figure 3:**
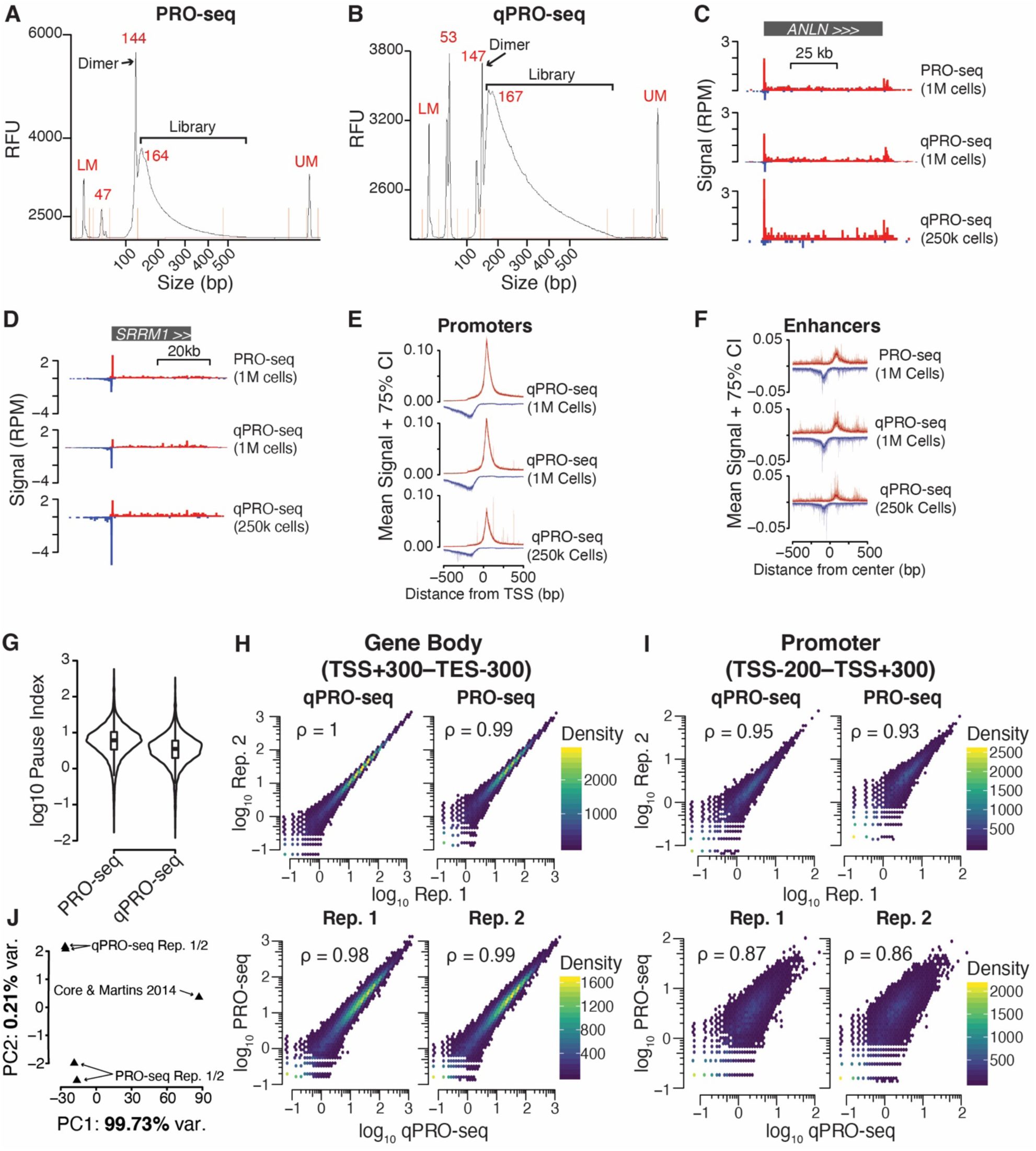
Direct comparison of PRO-seq and qPRO-seq in K562 cells. Bioanalyzer traces of libraries made using PRO-seq **(A)** and qPRO-seq. Library and adapter dimer are highlighed. **(B)**. Browser shots showing reads-per-million (RPM) normalized signal for PRO-seq (1M cells), qPRO-seq (1M cells), and qPRO-seq (250k cells) at *ANLN* **(C)** and *SRRM1* **(D)**. Metaprofiles of signal at promoters **(E)** and enhancers **(F)**. Mean and 75% confidence intervals are derived from 1000 subsamplings of 10% of regions. Enhancers are dREG peaks > 10 kb from a promoter. **(G)** Distribution of pause indicies at all genes in PRO-seq compared to qPRO-seq. **(H)** Correlation of RPM in gene body regions (TSS+300 to TES-300) between replicates and between protocols. **(I)** Correlation of RPM in promoter regions (TSS-200 to TSS+300) between replicates and between protocols. **(J)** Principal component analysis of PRO-seq, qPRO-seq, and published PRO-seq (Core et al. 2014) data in K562 cells. Variance explained (%) by each component is noted on the axis.

We have found that 10^6^ cells are sufficient input material for this new protocol, and that it can even be performed with as few as 0.05–0.25 x 10^6^ cells (Fig 3C–F). Data quality typically increases up to 10^6^ cells but much smaller improvements are seen by further increasing cell number. Polymerases are sampled from fewer positions overall in libraries made with 0.25 x 10^6^ cells, which causes data to look “spiky” and inflates read counts at highly expressed genes when normalizing per million mapped reads (Fig. 3C–F). Cell numbers required for high quality library generation will vary with the transcription activity of each cell type, with fast-dividing cultured cells typically showing the greatest activity.

We performed the new qPRO-seq protocol alongside the original protocol in two biological replicates of 10^6^ K562 cells (Fig. 3). We observe that the C1 beads decrease the bias against long RNAs seen with M280 beads, resulting in a relatively higher capture rate of gene body reads which results in lower overall pause indices (pause index is pause region signal divided by length-normalized gene body signal, so increased gene-body capture rate and unchanged pause region capture rate decreases pause index; Fig. 3G). In aggregate profiles, we observe no detectable difference in promoter or enhancer profiles across protocols (Fig. 3E–F). Though the overall number of unique reads obtained are lower in libraries made with 0.25 x 10^6^ cells as is expected with scarce input material, we still observe transcription at many enhancers (Fig. 3F). We find that both assays can reproducibly quantify both gene body and promoter-proximal polymerases (Fig. 3H–I) and discover similar sets of regulatory elements using dREG (Wang et al., 2019). The promoter region is more inherently variable, but using principal component analysis, we were able to determine that when compared to published, deeply sequenced PRO-seq data in K562 cells (Core et al., 2014), batch effect and/or sequencing depth accounted for >99% of variance in both the gene body and promoter region, while choice of protocol (PRO-seq or qPRO-seq) accounted for less than 0.3% of variance (Fig. 3J). This improved and optimized protocol expands the utility and accessibly of PRO-seq without sacrificing precision or data quality.

## Data Availability

K562 PRO-seq and qPRO-seq raw sequencing data, processed bigWig files and dREG peaks files have been deposited in GEO (GSE150625).

## Code Availability

Code used to analyze data and generate figures in this manuscript is available at https://github.com/JAJ256/qPRO. The pipeline used for PRO-seq alignment and processing is available at https://github.com/JAJ256/PROseq_alignment.sh. “Browser shots” were generated using code found here: https://github.com/JAJ256/browser_plot.R.

## Author Contributions

J.J. performed PRO-seq and qPRO-seq experiments and analyzed data. J.J., L.J.C., L.M.W, and L.A.W wrote and revised the manuscript. L.A.W., L.M.W., G.J.V., and A.D. optimized on-bead steps biochemical steps. All authors reviewed and approved the final manuscript. E.J.R. contributed the qPCR protocol. E.M.W. performed experiments to optimize the protocol for low-input experiments. All authors contributed experimentally or conceptually to the development of this protocol.

## Protocol Availability

The protocol presented in this manuscript is also available at https://dx.doi.org/10.17504/protocols.io.57dg9i6 and will be updated as further improvements are made.

## Acknowledgements

We thank the Cornell BRC Genomics facility and Peter Schweitzer for assistance with Illumina sequencing. We acknowledge many colleagues who have now used PRO-seq for various applications that we did not cite in this manuscript due to space constraints. This work was supported by NIH grants R01-GM025232 (to J.T.L.), R21-HG009021 and R35-GM128857 (to L.J.C.), and R01-HG009309 (to C.G.D). J.J. was supported by NHGRI fellowship F31-HG010820. The content is solely the responsibility of the authors and does not necessarily represent the official views of the National Institutes of Health.

## 2. Materials

! **CRITICAL:** Care should be taken to avoid nuclease contamination. Change gloves routinely and prepare/use nuclease-free reagents.

### 2.1 Chemicals and Reagents

1. Diethyl pyrocarbonate (DEPC; Sigma-Aldrich, cat. no. D5758) ! **CAUTION:** DEPC is toxic and harmful
2. DEPC treated ddH_2_O (0.1% v/v)
3. 4 M KCl (see Note 1)
4. 5 M NaCl (see Note 1)
5. 1M MgCl_2_ (see Note 1)
6. 10% (v/v) Triton X-100 (see Note 2)
7. 2% (v/v) Sarkosyl (Sigma-Aldrich, cat. no. L5125) (see Note 2)
8. 2% (v/v) Tween-20 (see Note 2)
9. 10 % Igepal® CA-630 (Millipore Sigma cat. no. I8896) (see Note 2)
10. 0.1 M DTT (see Note 2)
11. 1 M Sucrose (see Note 2)
12. 0.5 M EDTA, pH 8.0 (see Note 2)
13. 0.1 M EGTA, pH 8.0 (see Note 2)
14. Tris-Cl, pH 6.8 (See Note 2)
15. Tris-Cl, pH 7.4 (See Note 2)
16. Tris-Cl, pH 8.0 (See Note 2)
17. Glycerol
18. 1 N NaOH
19. 100% Ethanol
20. 75% Ethanol
21. Biotin-11-CTP 10 mM (PerkinElmer, cat. no. NEL542001EA)
22. Biotin-11-UTP 10 mM (PerkinElmer, cat. no. NEL543001EA)
23. Biotin-11-GTP 10 mM (PerkinElmer, cat. no. NEL545001EA)
24. Biotin-11-ATP 10 mM (PerkinElmer, cat. no. NEL544001EA)
25. GTP, 100 mM (Roche, cat. no. 11277057001)
26. ATP, 100 mM (Roche, cat. no. 11277057001)
27. Trypan Blue
28. Dynabeads™ MyOne™ Streptavidin C1 Beads (Invitrogen, cat. no. 65002)
29. TRIzol™ Reagent (Invitrogen, cat. no. 15596018)
30. TRIzol™ LS Reagent (Invitrogen, cat. no. 10296028) (optional, see Note 3)
31. Total RNA Purification Kit (Norgen, cat. no. 37500) (optional, see Note 3)
32. Chloroform
33. GlycoBlue™ (Invitrogen, cat. no. AM9515)
34. SUPERase-In™ RNase Inhibitor (20 U/µL) (Invitrogen, cat. no. AM2694)
35. Pierce™ Protease Inhibitor Tablets (Thermo Scientific cat. no. A32963)
36. T4 RNA Ligase (10 U/µL), supplied with 10X T4 RNA ligase buffer, 10 mM ATP and 50% (w/v) PEG8000 (NEB cat. no. M0204S)
37. RNA 5’ pyrophosphohydrolase (RppH; 5 U/µL) (NEB, cat. no. M0356S)
38. 10X ThermoPol Reaction Buffer (NEB, cat. no. B9004S)
39. T4 polynucleotide kinase (PNK; 10 U/µL), supplied with 10X PNK buffer (NEB, cat. no. M0201)
40. Maxima H Minus Reverse Transcriptase (200 U/µL) (Thermo Scientific, cat. no. EP0751)
41. dNTP mix, 12.5mM each (Roche, cat. no. 03622614001)
42. Q5® High-Fidelity DNA Polymerase, supplied with 5X Q5 Reaction Buffer and 5X High GC-Enhancer (NEB cat. no. M0491S)
43. Micro Bio-Spin™ RNase free P-30 Gel Columns (Bio-Rad, cat. no. 7326250) (optional, see Note 3)
44. RNA and DNA oligos (IDT DNA) (see Table 1 and Notes 4–6)
45. Agencourt AMPure XP SPRI beads (Beckman-Coulter, cat. no. A63880)
46. SYBR Gold stain (Invitrogen, cat. no. S11494)
47. Costar® Spin-X® 0.22 µm Centrifuge Tube Filters (Corning, cat. no. 1860)
48. SsoAdvanced Universal SYBR Green Supermix (BioRad, cat. no. 1725270)

**Table 1.**
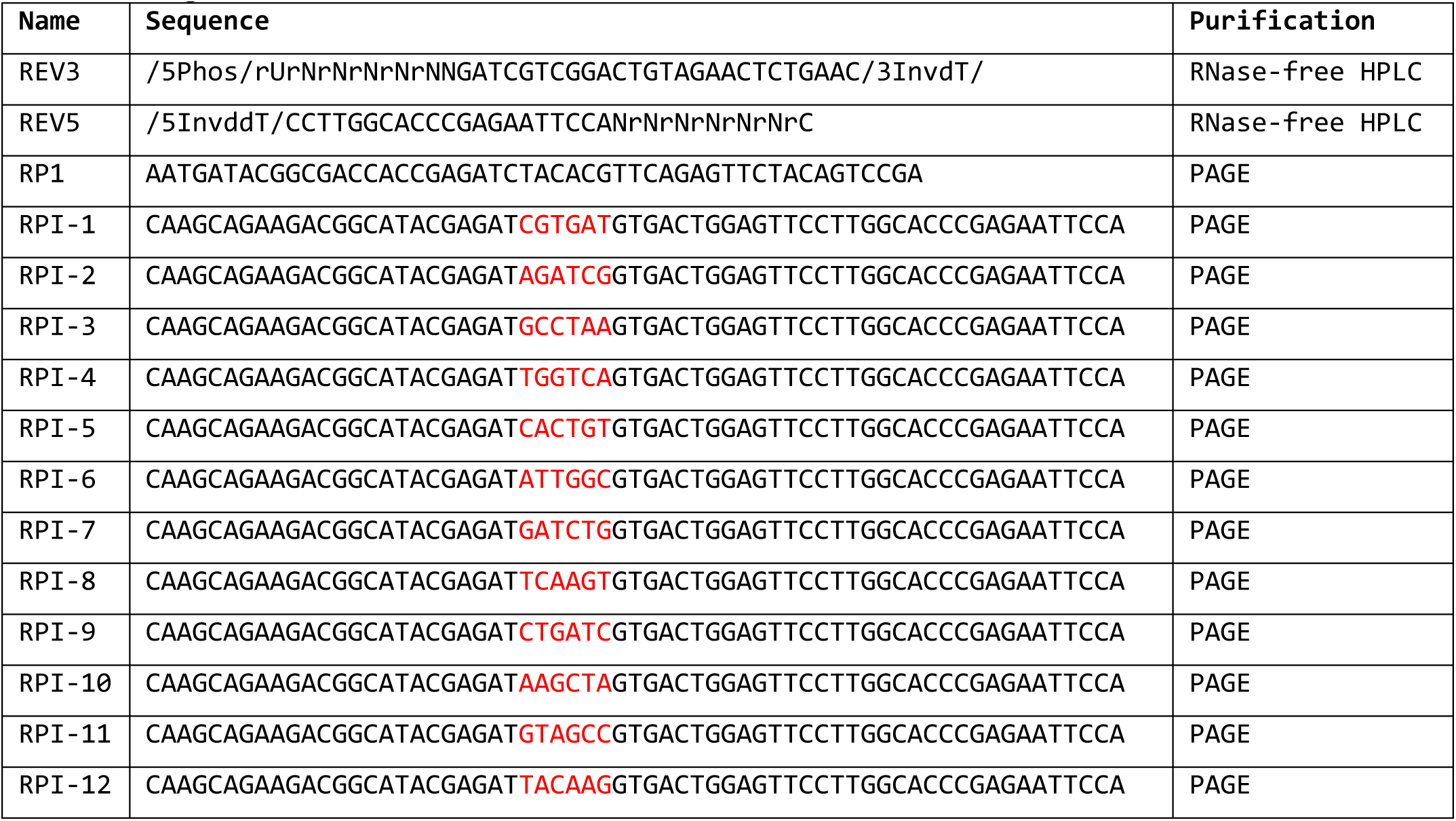
Oligonucleotides.

### 2.2 Buffers

1. <u>Permeabilization Buffer:</u> 10 mM Tris-Cl [pH 8.0], 10 mM KCl, 250 mM Sucrose, 5 mM MgCl_2_, 1 mM EGTA, 0.1% (v/v) Igepal, 0.5 mM DTT, 0.05% (v/v) Tween-20, and 10% (v/v) Glycerol in DEPC H_2_O. Add 1 Pierce protease inhibitor tablet and 10 µl SUPERase-In RNase inhibitor per 50 mL (see Note 7)
2. <u>Cell Wash buffer:</u> 10 mM Tris-Cl [pH 8.0], 10 mM KCl, 250 mM sucrose, 5 mM MgCl_2_, 1 mM EGTA, 0.5 mM DTT, and 10% (v/v) Glycerol in DEPC H_2_O. Add 1 Pierce protease Inhibitor tablet and 10 µl SUPERase-In RNase inhibitor per 50 mL (see Note 7)
3. <u>Freeze buffer:</u> 50 mM Tris-Cl [pH 8.0], 40% (v/v) glycerol, 5 mM MgCl_2_, 1.1 mM EDTA, and 0.5 mM DTT in DEPC H_2_O. Add 10 µl SUPERase-In RNase inhibitor per 50 mL (see Note 7)
4. <u>Bead Preparation Buffer:</u> 0.1 N NaOH and 50 mM NaCl in DEPC H_2_O.
5. <u>Bead Binding Buffer:</u> 10 mM Tris-HCl [pH 7.4], 300 mM NaCl, 0.1% (v/v) Triton X-100, 1 mM EDTA in DEPC H_2_O. Add 2 µl SUPERase-In Rnase Inhibitor per 10 mL (see Note 7)
6. <u>High Salt Wash buffer:</u> 50 mM Tris-HCl [pH 7.4], 2 M NaCl, 0.5% (v/v) Triton X-100, and 1 mM EDTA in DEPC H_2_O. Add 2 µl SUPERase-In RNase Inhibitor per 10 mL (see Note 7)
7. <u>Low Salt Wash Buffer:</u> 5 mM Tris-HCl [pH 7.4], 0.1% (v/v) Triton X-100, and 1 mM EDTA in DEPC H_2_O. Add 2 µl SUPERase-In RNase Inhibitor per 10 mL (see Note 7)
8. <u>2X Run-On Master Mix (2XROMM):</u> 10 mM Tris-Cl [pH 8.0], 5 mM MgCl_2_, 1 mM DTT, 300 mM KCl, 40 µM Biotin-11-CTP, 40 µM Biotin-11-UTP, 40 µM Biotin-11-ATP, 40 µM Biotin-11-GTP, 1% (w/v) Sarkosyl in DEPC H_2_O. Add 1 µL SUPERase-In RNase Inhibitor per reaction (see Note 8).

### 2.3 Consumables

1. Low-bind, nuclease-free microcentrifuge tubes (0.5 and 1.5 mL)
2. Low-bind, nuclease-free, filtered pipette tips
3. Wide bore, low-bind, nuclease-free, filtered P1000 and P200 tips (or cut standard bore ∼1cm from tip)

## 3. Methods

### 3.1 Cell Permeabilization

! **CRITICAL:** ALL steps should be carried out on ice or in a cold room

1. Prepare permeabilization buffer, wash buffer, and freeze buffer and place on ice (see Note 7).
2. Proceed using one of the following options: Option 3.1.2.1: Adherent cells (volumes are for 10 cm plates):
  1. Wash cells with 10 mL ice cold PBS.
  2. Repeat the PBS wash step for a total of two washes.
  3. Add 5 mL ice cold permeabilization buffer, scrape cells, and transfer to a conical tube.
  4. Rinse plate with 5 mL permeabilization buffer and pool cells in conical tube (V_f_ = 10 mL).
  Option 3.1.2.2: Suspension cells:
  1. Transfer cells into conical tube and spin down at 700–1000 x g for 4 min at 4°C (see Notes 9–10).
  2. Wash with 10 mL ice cold PBS (see Notes 10–11).
  3. Repeat the PBS wash for a total of two washes.
  4. Resuspend in 10 mL cold permeabilization buffer (see Note 11).
3. Continue here from step 3.1.2.1.4 or 3.1.2.2.4.
4. Incubate on ice for 5 min.
5. Check for permeabilization with Trypan blue. Greater than 98% permeabilization is ideal (see Note 12).
6. Spin down at 700–1000 x g for 4 min at 4°C (see Notes 9–10).
7. Wash with 10 mL ice cold cell wash buffer (see Notes 10–11).
8. Repeat the cell wash buffer wash for a total of two washes.
9. Decant wash buffer, and then carefully pipette off remaining buffer and discard without disturbing the cell pellet.
10. Using wide-bore tips, resuspend in 250 µL cold freeze buffer and transfer to a 1.5 mL tube.
11. Rinse the conical tube with an additional 250 µL freeze buffer and pool (V_f_ = 500 µL).
12. Count cells and add permeabilized spike-in cells if desired (see Notes 13– 14).
13. Spin down at 1000 x g for 5 min at 4°C (see Note 15).
14. Resuspend the desired number of cells for each run-on reaction in 52 µL freeze buffer (see Note 16).
15. Continue to the run-on or snap freeze 52 µL aliquots in LN_2_ and store at - 80°C (see Note 17).

### 3.2 Preparation

1. Pre-chill a microcentrifuge to 4°C.
2. Set a heat block with water in the wells to 37°C and another to 65°C and allow temperature to equilibrate.
3. Prepare bead preparation buffer, high salt wash buffer, low salt wash buffer, and binding buffer (see Note 7).
4. For each run-on reaction, wash 10 μL Dynabeads™ MyOne™ Streptavidin C1 Beads once in 1 mL bead preparation buffer using a magnet stand. Beads can be washed in bulk (see Notes 18–19).
5. Wash beads twice with 1 mL binding buffer (see Note 19–20).
6. Resuspend the beads in 25 μL binding buffer per sample. Place beads on ice or at 4°C until needed.

### 3.3 Run-On Reaction

1. Prepare 2XROMM equilibrate at 37°C (30°C for Drosophila) (see Note 8 and 21).
2. Using a wide bore tip, add 50 µL of permeabilized cells to new 1.5 mL tube.
3. Pipette 50 µL of preheated 2XROMM into each reaction tube (already containing permeabilized cells). *<u>Gently and thoroughly pipette the mixture 15 times</u>*. It is extremely important to thoroughly mix the reaction so that nucleotides diffuse into highly viscous chromatin!
4. Incubate in a thermomixer at 37°C (30°C for Drosophila) at 750 RPM for 5 min. Have RL buffer from Norgen kit or TRIzol LS ready for use.
5. Proceed to step 3.4.1.1 or 3.4.1.2 depending on choice of RNA extraction method immediately after the 5 min reaction is complete (take the sample off the heat block and immediately add buffer RL or TRIzol LS).

Use the table below to coordinate timing of run-on reaction and addition of TRIzol or Buffer RL. Start the timer (counting up) when you add the first sample:

| Time | Step |
| --- | --- |
| 00:00 | Mix permeabilized cells with 2XROMM for Sample 1 |
| 01:00 | Mix permeabilized cells with 2XROMM for Sample 2 |
| 02:00 | Mix permeabilized cells with 2XROMM for Sample 3 |
| 03:00 | Mix permeabilized cells with 2XROMM for Sample 4 |
| 04:00 | Mix permeabilized cells with 2XROMM for Sample 5 |
| 05:00 | Add buffer RL or TRIzol LS to Sample 1, vortex briefly |
| 06:00 | Add Buffer RL or TRIzol LS to Sample 2, vortex briefly |
| 07:00 | Add Buffer RL or TRIzol LS to Sample 3, vortex briefly |
| 08:00 | Add Buffer RL or TRIzol LS to Sample 4, vortex briefly |
| 09:00 | Add Buffer RL or TRIzol LS to Sample 5, vortex briefly |

### 3.4 Total RNA Extraction and Base Hydrolysis

1. Proceed from step 3.3.5 to one of the following options: Option 3.4.1.1: NORGEN RNA Extraction (see Note 3):
  1. Add 350 µL RL buffer and vortex.
  2. Add 240 µL 100% ethanol and vortex.
  3. Apply solution to Norgen RNA extraction column.
  4. Spin at 3,500 x g for 1 min at 25°C.
  5. Add 400 µL wash solution A (ensure ethanol has been added).
  6. Spin at 14,000 x g for 1 min at 25°C.
  7. Discard flow through.
  8. Repeat wash (steps 6 & 7) for a total of two washes.
  9. Spin at 14,000 x g for 2 min to dry column.
  10. Add 50 µL DEPC H_2_O and vortex.
  11. Elute by spinning at 200 x g for 2 min at 25°C and then at 14,000 x g for 1 min at 25°C.
  12. Elute again with 50 µL H_2_O and pool eluates (V_f_ = 100 µL).
  13. Denature at 65°C for 30 sec and then snap cool on ice.
  14. Add 25 *μ*L ice cold 1 N NaOH and incubate 10 min on ice.
  15. Add 125 *μ*L cold 1 M Tris-Cl pH 6.8, mix by pipetting.
  16. Add 5 µL 5 M NaCl and 1 µL GlycoBlue and mix.
  17. Add 625 µL 100% Ethanol and vortex (see Note 22).
  18. Centrifuge the samples at >20,000 x g for 20 minutes at 4°C (see Note 23).
  19. Carefully pipette supernatant off and discard (see Note 24).
  20. Add 750 µL 70% ethanol.
  21. Mix by gentle inversion and spin down briefly.
  22. Carefully pipette supernatant off and discard (see Note 24).
  23. Airdry the RNA pellet (see Note 25).
  24. Resuspend in 6 µL DEPC H_2_O.

Option 3.4.1.2: Trizol LS RNA Extraction (See Note 3):

1. Add 250 µL TRIzol LS with a wide bore P1000 tip, pipette vigorously but carefully >10X until all white globs of nucleoproteins are homogenized.
2. Pipette mix again with a standard bore P1000 tip. Samples should be completely homogenous.
3. Vortex vigorously for at least 15 seconds.
4. Incubate samples on ice until all run-on reactions are complete.
5. Add 65 µL chloroform (see Note 26).
6. Vortex the samples at max speed for 15 sec, then incubate on ice for 3 min.
7. Centrifuge the samples at >20,000 x g for 8 min at 4°C.
8. Transfer the ∼200 µL aqueous phase into a new tube (see Note 27).
9. Add 1 µL of GlycoBlue and mix.
10. Add 2.5X volumes (∼500 µL) 100% ethanol and vortex.
11. Centrifuge at >20,000 x g for 20 min at 4°C (see Note 23).
12. Carefully pipette supernatant off and discard (see Note 24).
13. Add 750 µL 70% ethanol.
14. Mix by gentle inversion and quickly spin down.
15. Carefully pipette supernatant off and discard (see Note 24).
16. Airdry the RNA pellet (see Note 25).
17. Resuspend in 30 µL DEPC H_2_O.
18. Briefly denature at 65°C for 30 sec and then snap cool on ice.
19. Add 7.5 *μ*L ice cold 1 N NaOH and incubate on ice for 10 min.
20. Add 37.5 *μ*L 1 M TrisCl pH 6.8, mix by pipetting.
21. Pass through a calibrated Bio-Rad RNase free P-30 column (follow manufacturer’s instructions).
22. Bring volume to 200 µL with DEPC H_2_O (add ∼ 125 µL).
23. Add 1 µL Glycoblue and 8 µL 5 M NaCl and vortex.
24. Add 500 µL 100% ethanol and vortex (see Note 22).
25. Centrifuge at >20,000 x g for 20 minutes at 4°C (see Note 23).
26. Carefully pipette supernatant off and discard (see Note 24).
27. Add 750 µL 70% ethanol.
28. Mix by gentle inversion and quickly spin down.
29. Carefully pipette supernatant off and discard (see Note 24).
30. Airdry the RNA pellet (see Note 25).
31. Resuspend in 6 µL DEPC H_2_O.

### 3.5 3’ RNA Adaptor Ligation

1. Continue here from step 3.4.1.1.24 or 3.4.1.2.31:
2. Add 1 µL 10 µM VRA3 (see Note 28).
3. Denature at 65°C for 30 sec and snap cool on ice.
4. Prepare ligation mix in the following order (see Note 29):

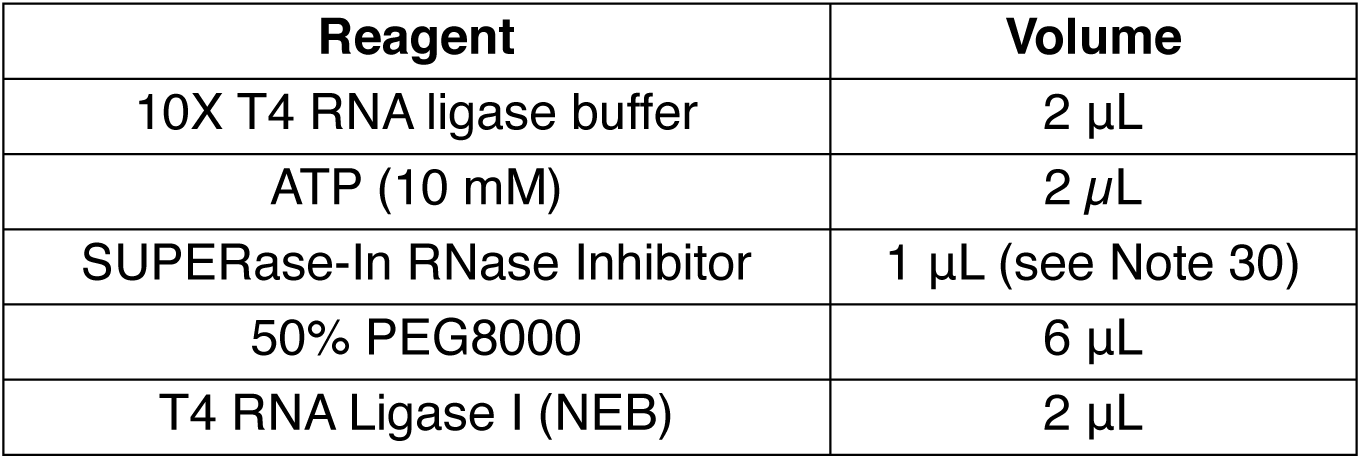
5. Add 13 *μ*L and mix by pipetting 10–15X (V_f_ = 20 µL).
6. Incubate at 25°C for 1 h.

### 3.6 Streptavidin Bead Binding

1. Add 55 µL binding buffer to each sample (V_f_ = 75 µL).
2. Add 25 *μ*L pre-washed beads to each sample (V_f_ = 100 µL).
3. Incubate for 20 min at 25°C with end to end rotation.
4. Wash once with 500 *μ*L High Salt Wash buffer with tube swap (see Notes 19, 31–32).
5. Wash once with 500 *μ*L Low Salt Wash buffer (see Notes 19, 31, 33).

### 3.7 On-Bead 5’ Hydroxyl Repair

1. Resuspend beads in 19µL PNK mix (V_f_ = 20 µL; see Note 34):

2. Incubate at 37°C for 30 min (see note 35).

| Reagent | Volume |
| --- | --- |
| DEPC H<sub>2</sub>O | 13 μL |
| 10X PNK buffer | 2 μL |
| 10 mM ATP | 2 μL |
| T4 PNK (NEB) | 1 μL |
| SUPERase-IN RNase Inhibitor | 1 μL (see Note 30) |

### 3.8 On-Bead 5’ Decapping

1. Place the tubes on a magnet stand and remove supernatant (see Notes 19– 20, 33).
2. Resuspend the beads in 19 µL RppH mix (V_f_ = 20 µL; see Notes 34, 36):

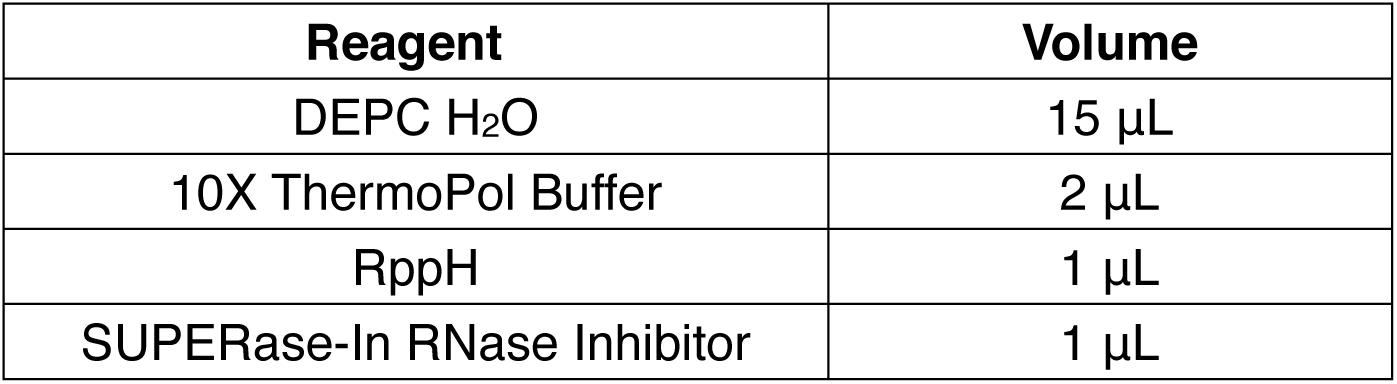
3. Incubate at 37°C for 1 h (see note 35).

| Reagent | Volume |
| --- | --- |
| DEPC H<sub>2</sub>O | 15 μL |
| 10X ThermoPol Buffer | 2 μL |
| RppH | 1 μL |
| SUPERase-In RNase Inhibitor | 1 μL |

### 3.9 On-Bead 5’ RNA Adaptor Ligation

1. Place the tubes on a magnet stand and remove supernatant (see Notes 19– 20, 33).
2. Resuspend the beads in 7 µL adapter mix (V_f_ = 8 µL):

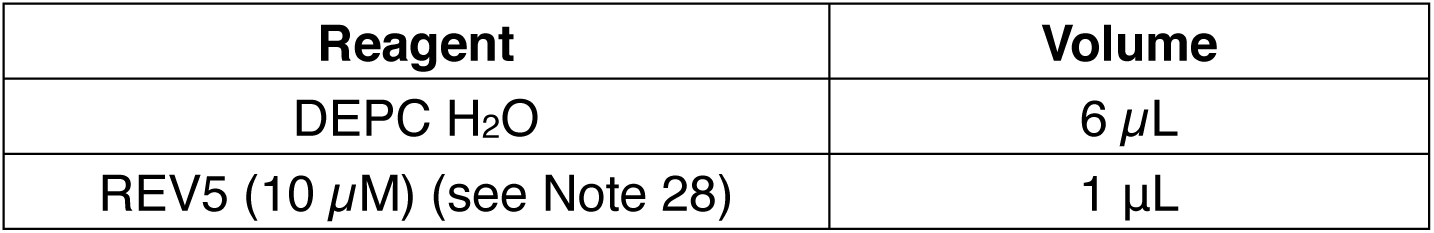
3. Denature at 65°C for 30 sec, then snap cool on ice.
4. Prepare ligation mix in the following order (see Note 29):

| Reagent | Volume |
| --- | --- |
| DEPC H<sub>2</sub>O | 6 μL |
| REV5 (10 μM) (see Note 28) | 1 μL |

5. Add 12 µL to each tube (V_f_ = 20 µL).
6. Incubate at 25°C for 1 h (see Note 35).

3. Denature at 65°C for 30 sec, then snap cool on ice. 4. Prepare ligation mix in the following order (see Note 29):
| Reagent | Volume |
| --- | --- |
| 10x T4 RNA ligase buffer | 2 μL |
| ATP (10 mM) | 2 μL |
| SUPERase-In RNase Inhibitor | 1 μL (see Note 30) |
| 50% PEG8000 | 6 μL |
| T4 RNA ligase I | 1 μL |

### 3.10 TRIzol Elution of RNA

1. Wash once with 500 μL High Salt Wash buffer with tube swap (see Notes 19, 31–32).
2. Wash once with 500 μL Low Salt Wash buffer (see Notes 19, 31, 33).
3. Resuspend beads in 300 *µ*L TRIzol.
4. Vortex at max speed for >20 sec, then incubate on ice for 3 min.
5. Add 60 *µ*L chloroform (see Note 26).
6. Vortex at max speed for 15 sec, then incubate on ice for 3 min.
7. Centrifuge at >20,000 x g for 8 min at 4°C.
8. Transfer the aqueous phase (∼180 *µ*L) to a new tube (see Note 27).
9. Add 1 *µ*L GlycoBlue and mix.
10. Add 2.5X volumes (∼450 *μ*L) 100% ethanol and vortex.
11. Centrifuge the samples at >20,000 x g for 20 min at 4°C (see Note 23).
12. Carefully pipette supernatant off and discard (see Note 24).
13. Add 750 *μ*L 70% ethanol.
14. Mix by gentle inversion and quickly spin down.
15. Carefully pipette supernatant off and discard (see Note 24).
16. Airdry the RNA pellet (see Note 25).

### 3.11 Off-Bead Reverse Transcription

1. Resuspend RNA pellet in 13.5 *μ*L RT resuspension mix (see Note 37):

2. Denature at 65°C for 5 min and snap cool on ice.
3. Prepare RT master mix:

| Reagent | Volume |
| --- | --- |
| DEPC H<sub>2</sub>O | 8.7 μL |
| RP1 (10 μM) | 4 |
| dNTP mix (12.5 mM each) | 0.8 μL |

4. Add 6.5 *μ*L to each sample (V_f_ = 20 *μ*L).
5. Cycle as follows: 50°C for 30 min, 65°C for 15 min, 85°C for 5 min, then hold at 4°C.
6. Immediately proceed to PreCR, test amplification, or full-scale amplification. Samples can be stored overnight at -20°C (see Notes 38–39).

2. Denature at 65°C for 5 min and snap cool on ice. 3. Prepare RT master mix:
| Reagent | Volume |
| --- | --- |
| 5X RT Buffer | 4 μL |
| 100 mM DTT | 1 μL |
| SUPERase-In RNase Inhibitor | 0.5 μL (see Note 30) |
| Maxima H Minus RT enzyme | 1 μL |

### 3.12 PreCR (Optional, see Note 38)

1. Add 2.5 *μ*L RPI-n indexed primer (10 µM) to each sample. Use different barcodes for samples that will be pooled and sequenced together.
2. Prepare the PreCR master mix:

3. Add 77.5 μL of the PreCR mix to each sample for final volume 100 *μ*L (see Note 40).
4. Amplify libraries for 5 cycles on thermal cycler using the following settings:
  a. 95°C for 2 min
  b. 95°C for 30 sec
  c. 56°C for 30 sec
  d. 72°C for 30 sec
  e. Go to step 2 for 4 more times
  f. 72°C for 5 min
  g. Hold at 4°C.
5. Store samples at -20°C or proceed to test amplification.

| Reagent | Volume |
| --- | --- |
| ddH<sub>2</sub>O | 33.5 μL |
| 5x Q5 buffer | 20 μL |
| 5x Q5 enhancer | 20 μL |
| Primer RP1 (10 μM) | 1 μL |
| dNTP mix (12.5 mM each) | 2 μL |
| Q5 polymerase (NEB) | 1 μL |

### 3.13 Test Amplification (Gel) (Optional, see Note 39)

1. Make the first dilution:
  a. If PreCR was performed, add 7.7 μL of the 100 *μ*L PreCR reaction to 0.3 *μ*L ddH_2_O for a final volume of 8 *μ*L (see Note 41).
  b. If PreCR was skipped, add 1.54 *μ*L of the 20 *μ*L RT reaction to 6.46 *μ*L ddH_2_O for a final volume of 8 *μ*L (see Note 41).
2. Make 4-fold serial dilutions by adding 2 μL of each dilution to 6 μL ddH_2_O for the next dilution.
3. Remove and discard 2 *μ*L from the final dilution (all dilutions should now be 6 *μ*L).
4. Choose a target number of total cycles for test amplification using the table below (see Note 42). The first dilution simulates full-scale amplification at the total number of cycles (PreCR cycles + Test Amp cycles) minus 4. Subtract 2 cycles sequentially for the following dilutions.

5. Make test PCR mix:

| Dilution | 1 | 2 | 3 | 4 | 5 | 6 | 7 | 8 |
| --- | --- | --- | --- | --- | --- | --- | --- | --- |
| 19 Total Cycles | 15 | 13 | 11 | 9 | 7 | 5 | 3 | 1 |
| 21 Total Cycles | 17 | 15 | 13 | 11 | 9 | 7 | 5 | 3 |
| 23 Total Cycles | 19 | 17 | 15 | 13 | 11 | 9 | 7 | 5 |

6. Add 14μL PCR mix to the 6 *μ*L diluted test samples (V_f_ = 20 *μ*L).
7. Amplify reactions for the desired amount of cycles using following settings: ! **CRITICAL** Remember to account for PreCR. Subtract 5 cycles from your total target test amplification cycles.
  a. 95°C for 2 min
  b. 95°C for 30 sec
  c. 65°C for 30 sec (see Note 43)
  d. 72°C for 30 sec
  e. Go to step 2 for the desired number of cycles
  f. 72°C for 5 min
  g. Hold at 4°C
8. Mix with gel loading dye to 1X and run 10 *μ*L on a 2.2% Agarose gel or run 2 *μ*L on a native 8% polyacrylamide gel and stain with SYBR Gold.
9. Use the test amplification gel to determine the appropriate number of cycles for full-scale amplification (see Note 44).

5. Make test PCR mix:
| Reagent | Volume |
| --- | --- |
| H<sub>2</sub>O | 4.4 µL |
| 5x Q5 buffer | 4 µL |
| 5x Q5 enhancer | 4 µL |
| Primer RP1 (10 µM) | 0.5 µL |
| Primer RPI-n (10 µM) | 0.5 µL |
| dNTP mix (12.5 mM each) | 0.4 µL |
| Q5 polymerase (NEB) | 0.2 µL |

### 3.14 Test Amplification (qPCR) (Optional, see Note 39)

1. Add 1.54 *μ*L of the 20 *μ*L RT reaction to 0.46 *μ*L ddH_2_O (V_f_ = 2 *μ*L).
2. Make the qPCR master mix:

3. Add 8 *μ*L of the qPCR master mix to 2 *μ*L diluted RT reaction (V_f_ = 10 *μ*L).
4. Quickly spin plate to collect liquid.
5. Amplify in a real-time PCR system using the following conditions:
  a. Amplification
    i. 98 °C for 2 min
    ii. 98 °C for 15 sec
    iii. 60 °C for 60 sec
    iv. Go to step 2 for 39 additional cycles
  b. Melting Curve
    i. 95 °C for 15 sec
    ii. 60 °C for 1 min
    iii. 96 °C for 15 sec
    iv. 60 °C for 16 sec
6. Calculate the number of full-scale amplification cycles as the cycle number where Rn reaches 0.25 × Rn_max_.

| Reagent | Volume |
| --- | --- |
| Primer RP1 (10 μM) | 0.25 μL |
| Primer RPI-n (10 μM) | 0.25 μL |
| 2X SsoAdvanced Universal SYBR Green Supermix | 5 μL |
| H<sub>2</sub>O | 2.5 μL |

### 3.14 Full-Scale Amplification

1. If PreCR and Test Amplification were skipped:
  a. Add 2.5 *μ*L of an RPI-n indexed primer (10µM) to each 20 *μ*L RT reaction. Use different barcodes for samples that will be pooled and sequenced on a single lane.
  b. Prepare the PCR master mix in step 2 of section 3.12 (PreCR).
  c. Add 77.5 *μ*L PCR master mix to each sample for final volume 100 *μ*L (see Note 40).
  d. Run the desired number of cycles (see below)
2. If PreCR was skipped but Test Amplification was performed:
  a. Add 2.5 *μ*L of an RPI-n indexed primer (10µM) to the remaining 18.5 *μ*L RT reaction. Use different barcodes for samples that will be pooled and sequenced on a single lane.
  b. Prepare the PCR master mix in step 2 of section 3.12 (PreCR) but use 35 *μ*L ddH_2_O instead of 33.5 *μ*L.
  c. Add 79 *μ*L PCR master mix to each sample for final volume 100 *μ*L (see Note 40).
  d. Run the desired number of cycles (see below).
3. If PreCR and Test Amplification were performed:
  a. Prepare the following spike-in PCR mix:

  b. Add 7.7 *μ*L PCR spike-in mix to each sample for final volume 100 *μ*L (see Note 40).
  c. Run the desired number of cycles.
4. Amplify for the desired number of cycles: ! **CRITICAL:** Remember to account for PreCR. Subtract 5 cycles from your total target full-scale amplification cycles.
  a. 95°C for 2 min
  b. 95°C for 30 sec
  c. 65°C for 30 sec (see Note 43)
  d. 72°C for 30 sec
  e. Go to step 2 for the desired number of cycles
  f. 72°C for 5 min
  g. Hold at 4°C
5. Allow PCR reactions to reach room temperature.
6. Add 180 *μ*L SPRI beads (see Note 45) at room temperature and immediately mix by pipetting > 15X.
7. Incubate at room temperature for 5 min.
8. Place on a magnet stand and remove the supernatant.
9. Wash the beads twice with 70% Ethanol without resuspending. **! CRITICAL:** Do not disturb the beads or library recovery will be greatly reduced.
10. Airdry the beads for 5 min. Do not over dry the beads.
11. Resuspend beads in 22 *μ*L 10 mM Tris-Cl, pH 8.0 (no EDTA).
12. Incubate at room temperature for 5 min.
13. Place the beads on a magnet stand and transfer 20 *μ*L to a new tube.
14. Quantify the library using the Qubit dsDNA-HS assay and run on a Bioanalyzer.

| Reagent | Volume |
| --- | --- |
| ddH<sub>2</sub>O | 3.7 μL |
| 5x Q5 buffer | 1.5 μL |
| 5x Q5 enhancer | 1.5 μL |
| dNTP mix (12.5mM each) | 0.5 μL |
| Q5 polymerase (NEB) | 0.5 μL |

### 3.15 PAGE purification (optional, see Note 46)

1. Add Orange G loading dye to 1X to the entire library volume.
2. Run the samples on a native 8% polyacrylamide gel.
3. Stain with SYBR Gold.
4. Cut a gel slice from immediately above the adapter dimer to ∼650 bp (see Note 44).
5. Place the gel slice in a 0.5 mL microfuge tube.
6. Make a hole in the bottom of the tube with an 18G needle.
7. Nest the 0.5 mL tube in a 1.5 mL tube and spin at 5000 x g for 1 min.
8. If gel remains in the 0.5 mL tube, repeat step 7 and pool shredded gel fractions by suspending each in 250 *μ*L soaking buffer using a wide-bore P1000 tip.
9. Soak the gel pieces in 0.5 mL soaking buffer (TE + 150mM NaCl + 0.1% Tween-20) overnight with agitation at 37°C.
10. Spin the tube at 5000 x g for 1 min.
11. Pipette as much of the soaking buffer as possible without transferring gel pieces into a new tube.
12. Add an additional 0.5 mL soaking buffer and incubate 4 h at 37°C with agitation.
13. Spin the tube at 5000 x g for 1 min.
14. Pipette as much of the soaking buffer as possible without transferring gel pieces into the tube with the previous eluate.
15. Pass the remaining gel solution through a Costar Spin-X column using a cut P1000 tip and pool with the previous eluate (V_f_ = 1 mL)
16. Reduce the volume by half (V_f_ = 0.5 mL) using vacuum dryer at 37°C.
17. Add 1uL GlycoBlue.
18. Add 2.5X volume (1.25 mL) 100% ethanol and vortex.
19. Centrifuge at >20,000 x g for 20 minutes at 4°C (see Note 23).
20. Carefully pipette off the supernatant and discard (see Note 24).
21. Add 750µL of 75% ethanol.
22. Mix by gentle inversion and quickly spin down.
23. Carefully pipette off the supernatant and discard (see Note 24).
24. Air-dry the RNA pellet (see Note 25).
25. Resuspend the pellet in the desired volume of 10 mM Tris-Cl, pH 8.0 (no EDTA!).

## 4. Notes

1. All salt solutions should be prepared in ddH_2_O. Then add 0.1% (v/v) DEPC, stir overnight, and autoclave. Tris buffers instead need to be carefully prepared with DEPC-treated ddH_2_O.
2. All other solutions (detergents, DTT, sucrose, EDTA/EGTA, and Tris buffers) should be prepared in DEPC ddH_2_O in RNase free containers and filter sterilized. Glassware can be made RNase by filling with water, adding 0.1% (v/v) DEPC, incubating with agitation overnight, and autoclaving. Alternatively, glassware can be baked at 300 °C for 4 hours.
3. TRIzol LS or the Norgen Total RNA Purification Kit can be used to extract total RNA from the run-on reaction. Both options produce identical results. The Norgen kit is faster and less technically challenging to use, but more expensive. If TRIzol LS is used, Micro Bio-Spin™ RNase free P-30 Gel Columns are also needed to remove unincorporated biotin-NTPs as the biotin concentration will otherwise overwhelm the binding capacity of the streptavidin beads.
4. REV3 and REV5 sequences are reverse complements of the standard RA3/RA5 adapter design used in the Illumina TruSeq small RNA library prep kit with added 6XN UMIs and a final ligation-optimal fixed base. The oligos are blocked from 5. additional ligation by an inverted dT/ddT group. This design results in sequencing of the 3’ end of the nascent RNA at the beginning of read 1 using standard Illumina sequencing primers. Standard RA3/RA5 adapters (custom synthesized or from compatible library prep kits) can be substituted, but in this case paired end sequencing is necessary as the 3’ end of the nascent RNA will be sequenced at the beginning of read 2.
5. Index sequences in RPI-n primers are standard 6 nt Illumina TruSeq indices. If multiplexing of more than 12 libraries is desired, additional indexed primers can be designed using additional TruSeq 6 nt index sequences. The index sequences must be inserted in the primer reverse complemented because the RPI-n primers are the reverse primers in the library PCR reaction.
6. If desired, specialized REV3 oligos can be designed to facilitate additional in-line barcoding to allow for the pooling of multiple samples after the 3’ ligation, which simplifies downstream handling. We have had success with replacing the UMI sequence (rNrNrNrNNN) with known barcode sequences followed by the final fixed rU. These barcodes are then the first 6 sequenced nucleotides of Read 1 and can be used to computationally demultiplex samples.
7. The permeabilization buffer, cell wash buffer, freeze buffer, and bead washing/binding buffers can be made and filter-sterilized in advance without the DTT, SUPERase-In™ RNase Inhibitor, and Pierce™ protease inhibitor tablets. DTT, SUPERase-In™ RNase Inhibitor, and protease inhibitor tablets can be added when buffers are needed. Store buffers at 4°C. Use DEPC treated glassware or RNase free plasticware.
8. The run-on reaction uses 4 biotin-NTPs. However, ATP and GTP can be substituted at equal concentration for Biotin-11-ATP and Biotin-11-GTP to reduce cost. Biotin-11-ATP and biotin-11-GTP are 10X as expensive as biotin-11-CTP and biotin-11-UTP. With two biotin-NTPs blocking elongation, each polymerase can be expected to extend ∼5 nt or less which we find gives sufficient resolution for the vast majority of applications. For low cell number experiments, increase the concentration of the biotin-NTPs to 500 µM. Biotin-NTP incorporation efficiency is ∼60% with the concentration in the 2XROMM as written, which is sufficient for experiments using 10^6^ cells or greater, but increasing the concentration improves incorporation to ∼77% (data not shown).
9. Use a centrifuge with a swinging bucket rotor for all centrifuge steps during cell permeabilization. Using a fixed angle rotor will shear cells, releasing a smear of white chromatin.
10. Centrifuge speed is cell size dependent. We typically centrifuge HeLa at 800 x g and Drosophila at 1,000 x g.
11. When resuspending cells during permeabilization after centrifugation steps, first gently resuspend the cell pellet with 1 mL solution with a wide-bore P1000 tip. Then add the remaining volume (usually 9 mL) and mix by gentle inversion.
12. If your cell type is not permeabilized under these conditions, add Triton X-100 to 0.1-0.2%.
13. When processing multiple samples, if counting will cause the cells to sit on ice for greater than 10 min, reserve 10 *μ*L for counting, aliquot cells in 100 *μ*L aliquots, and snap freeze. Count the cells and then adjust the concentration with freeze buffer after thawing and prior to the run-on.
14. In order to robustly normalize between conditions where a dramatic change in global transcription levels are expected, we add a fixed number of cells of a different species to a fixed number of experimental cells at the permeabilization step. Reads can be mapped to a combined genome, and the number of spike-in mapped reads can then be used as a scaling factor. These cells should be permeabilized prior to the experiment, aliquoted, and added to 1-2% by cell number after permeabilization and counting, either just prior to freezing or just prior to the run-on reaction. We frequently use Drosophila S2 cells to normalize human cell experiments and vice versa.
15. Eppendorf tubes can be spun in a fixed angle rotor, but we continue to use a swinging bucket rotor so that cells collect at bottom of tube (this tends to decrease cell loss).
16. We have had success performing this protocol with as few as 50k primary human cells. In general, we find that the quality of libraries will increase until ∼1 × 10^6^ cells per run-on but using more cells than this offers little benefit. This will also depend on how transcriptionally active a given cell type is and genome size.
17. Permeabilized cells are stable indefinitely at -80°C (Chu et al., 2018).
18. C1 Streptavidin beads are preferred compared to M280 beads because they have higher binding capacity and use a negatively charged matrix. This significantly reduces carryover of non-biotinylated RNAs including adapter dimers.
19. Be careful not to disturb beads when removing buffers from tubes. Open tube caps prior to placing them on the magnet stand, as opening on the magnet stand can disturb the liquid. Check pipette tip against a white background before discarding liquid to ensure beads are not present.
20. Always quickly spin samples down using a picofuge to remove liquid from tube caps.
21. When preparing the 2XROMM, first add all components other than Sarkosyl and mix by vortexing on high for >10 sec. Collect the solution with a quick spin, add Sarkosyl, and mix thoroughly by pipetting carefully to avoid bubbles. If you leave the 2XROMM on ice, a precipitate can form. Before use, check if this has occurred. The precipitate can be re-dissolved by heating at 37 °C for ∼5 min and pipette mixing.
22. If the protocol needs to be performed over two days, the ethanol precipitation in 3.4 is the safest overnight stopping point. After step 17 of 3.4.1 or step 24 of section 3.4.2, samples can be stored at -80 °C. The protocol can also be stopped after the ethanol precipitation following TRIzol extractions, but samples must be stored for at most one night at 4 °C because at -20 °C precipitation of guanidinium salts can occur and interfere with enzymatic reactions. Alternatively, the RNA pellet can be stored at -80 °C after the precipitation and removal of the supernatant.
23. A blue pellet should be visible at the bottom of tube. The pellet can be difficult to see but should be visible. It may appear spread out. If a pellet is not visible, vortex well and repeat spin.
24. When removing the supernatant before the 70% ethanol wash be careful not to disturb the pellet. Approximately 30–50μL of ethanol can be left in the tube to avoid disturbing the pellet prior to adding the 70% ethanol wash. This procedure can also be used after the 70% ethanol wash, but then remove the final 30-50 *μ*L using a P200 tip after a quick spin in a picofuge.
25. Air dry the RNA pellet by leaving tubes open in fume hood to prevent contamination. This will take ∼3-10 min depending on how much ethanol is left in the tube. Do not to let the RNA pellet dry completely as this will greatly decrease its solubility.
26. When pipetting chloroform, always pipette twice because the first draw always leaks.
27. When transferring the aqueous phase of TRIzol extractions to a new tube, tilt the tube to a 45° angle and carefully remove only the clear liquid. Avoid contamination by the pink organic phase or white interphase.
28. The concentration of RNA adapters in the ligation steps (1 *μ*L 10 µM) is optimal for approximately 10^6^ mammalian cells. For lower cell numbers, the adapter concentration must be diluted to limit dimer formation. We dilute linearly with cell concentration relative to this established concentration, i.e. 1 *μ*L 5 µM for 5 × 10^5^ cells, 1 *μ*L 2.5 µM for 2.5 × 10^5^ cells, etc.
29. Pipette slowly because 50% PEG8000 is very viscous. Heating 50% PEG8000 makes it easier to pipette. Pipette the ligation mix until it is homogenous before use.
30. When preparing enzymatic reaction mixtures that contain SUPERase-In RNase Inhibitor, a fixed volume (1 *μ*L) SUPERase-In can be added to the entire master mix regardless of number of reactions to decrease cost. Bring the remainder of the master mix up to the required volume with DEPC H_2_O. Murine RNase inhibitor can also be substituted to limit costs for all steps after the run-on. SUPERase-In is recommended prior to the run-on as it inhibits T1 RNase.
31. For each washing step gently invert tubes 10–15X, quickly spin down with a picofuge, open caps, and then place on the magnet stand. Wait 1-2 minutes and pipette the supernatant off without disturbing the beads. If there are bubbles in the tube carefully pipette them off first and then remove supernatant. Beware that bubbles may dislodge beads from the side of the tube. After removing the bulk of the liquid, collect remaining liquid with a quick spin in a picofuge, place the tube back on the magnet stand, and carefully remove remaining liquid by pipetting.
32. Transferring beads to a new tube after the binding incubation—during the high salt wash step—helps limit adapter dimer formation. After resuspending the beads in High Salt buffer, quickly spin down with a picofuge, resuspend beads by gently pipetting, and carefully transfer to a new tube. Pipette slowly to avoid bead loss! Place this new tube on the magnet stand and proceed with the washing protocol.
33. Do not allow streptavidin beads to dry completely, as this can lead to clumping and make full resuspension impossible. When processing multiple samples, remove liquid from the previous wash or enzymatic step from the first sample and immediately resuspend those beads in the next solution, then repeat this process for additional samples.
34. On-bead reaction volumes assume that 1 *μ*L of liquid remains on the beads.
35. Mix on-bead reactions by gently flicking the tubes every 10 minutes.
36. We have also successfully used Cap-Clip™ Acid Pyrophosphatase (CELLTREAT) instead of RppH. Cap-Clip has lower buffer pH which may alleviate base hydrolysis of RNA that could occur in the pH 8.0 ThermoPol buffer. However, this is not a major concern except for in the most sensitive of applications.
37. Reverse transcription can also be performed on-bead, but we find that this significantly reduces library yield while increasing adapter dimer. For this reason, it is not recommended except in cases where material is abundant (10^7^ cells) and speed is paramount. To do this, follow steps 1–2 in section 3.12, then follow section 3.13, but resuspend the beads instead of the RNA pellet in RT resuspension mix. After RT, elute cDNA by heating the bead mixture to 95°C, quickly place tubes on a magnet stand, and remove and save supernatant. Resuspend beads in 20 *μ*L ddH_2_O and repeat the process for a final volume of 40 *μ*L. Proceed with PreCR but use 20 *μ*L less ddH_2_O (13.5 *μ*L) in the PreCR mix and use the entire 40 *μ*L eluate instead of the 20 *μ*L RT mix.
38. PreCR is optional if full scale amplification will be performed within 2 days. Longer storage of single-stranded cDNA libraries can lead to loss of library material. If you are skipping PreCR, simply store the 20 *μ*L RT reaction at -20°C overnight and perform test amplification the next day.
39. Because this protocol uses molecular barcodes (UMIs) which facilitate robust computational PCR deduplication, it is less important to precisely determine the optimal cycle number. We recommend performing test amplification the first time you perform this protocol with a given amount of material from a given cell line to determine the optimal cycle number. For future experiments where the material and cell number are constant, test amplification can be skipped. Adjust the volume of the full-scale PCR to 100 *μ*L total volume (accounting for the fact that the written protocol assumes loss due to test amplification). Test amplification can be performed either by PCR of a dilution curve and PAGE analysis or qPCR.
40. Do not attempt to scale down the PreCR or full-scale amplification steps to save PCR reagents. If RT reaction mixture exceeds 20% of the PCR reaction volume, significant inhibition of PCR will occur and lead to dramatically lower final library yield.
41. Taking 7.7 *μ*L of the 100 *μ*L PreCR reaction leaves 92.3 *μ*L for full-scale amplification. 25% of material in the first dilution is lost to make the next serial 4-fold dilution (2 of 8 *μ*L). Because (7.7 * 0.75) / 92.3 ≈ 1/16, this first dilution is equivalent to the number of test amplification cycles less 4. If starting from the RT reaction, the volume has been adjusted for 5-fold lower starting volume.
42. Additional cycles can vary by cell type. For HeLa, we typically perform 14 additional cycles (19 total cycles), which simulates 15 full-scale amplification cycles. For low input libraries (50k-250k mammalian cells), we typically perform 20 additional cycles (23 cycles total), which simulates 21 full-scale amplification cycles.
43. If PreCR was skipped, use an annealing temperature of 56°C for the first 5 cycles of test amplification and the full-scale amplification.
44. Desired amplification characteristics include a sufficient amount of product (smear starting ∼150 bp), no evidence of overamplification, and ∼50% primer exhaustion (Mahat et al., 2016). The adaptor dimer product is 132 bp, and the smear will start 15–20 bp above this band. RNA degradation will lead to shorter library products.
45. AMPure XP beads will work, as will any commercially available or homemade SPRI bead cleanup reagent based on PEG precipitation. Be sure to allow beads to reach room temperature, or excess primers will also precipitate.
46. Due to advances in streptavidin bead technology and titration of adapters presented in this protocol, PAGE purification is rarely necessary. We prefer to sequence libraries that are 0%–25% adapter dimer rather than risk size bias associated with gel purification. Only perform PAGE purification if absolutely necessary. If needed, multiple libraries can be pooled by molarity as determined by bioanalyzer and extracted from the same gel lane to minimize size bias.

